# Impact of Rigid Gas Permeable Contact Lenses Removal on Anterior and Posterior Corneal Surfaces in Keratoconus Patients

**DOI:** 10.1101/688929

**Authors:** Motohiro Itoi, Koji Kitazawa, Hisayo Higashihara, Chie Sotozono

**Author notes:** Correspondence author: Koji Kitazawa, (K.K). These authors contributed equally to this work.

## Abstract

**Purpose:** To evaluate the impact of removal of rigid gas-permeable (RGP) contact lenses on the anterior and posterior cornea surfaces of eyes with keratoconus.

**Methods:** Eight eyes of 8 patients with keratoconus (KC) (age 34.3 ± 15.3 years; range 19–60 years) were enrolled. Anterior segment optical coherence tomography (AS-OCT) was performed at 1, 5, 10, 20, and 60 minutes after the patients removed their RGP contact lenses. Measurements included anterior and posterior best-fit sphere (BFS); elevation values and corneal surface areas; corneal thickness at the thinnest point; and the anterior-posterior ratio of the corneal surface (As/Ps) between 1 minute and 60 minutes after RGP contact lens removal.

**Results:** Anterior and posterior elevation values and corneal surface areas showed significant increases, whereas anterior and posterior BFS and central corneal thickness decreased significantly (P < 0.01) between 1 minute and 60 minutes after RGP contact lens removal. No statistically significant differences were found in the As/Ps ratio during the first hour after suspending RGP contact lens wear.

**Conclusions:** We found that the patients with keratoconus experienced significant changes in both the anterior and posterior corneal shape for 60 minutes after removal of RGP contact lenses.

## Introduction

Keratoconus is an ectatic corneal disorder characterized by corneal steepening and apical corneal thinning [1]. As the disease progresses, eyeglasses can no longer provide adequate vison due to the irregular astigmatism created by ectasia of the central cornea. Consequently, keratoconus is primarily managed by wearing rigid gas-permeable (RGP) contact lenses [2], which change corneal astigmatism as a result of the optical characteristics of the tear layer behind the lens [3], and induce anterior and posterior topographic changes to the cornea [4-7].

Several studies have examined the topographic impact of suspending RGP contact lens wear on keratoconus in the eye and have demonstrated that the anterior shape of patient’s cornea adopts its truest and perhaps most irregular shape after RGP contact lens removal, as evidenced by keratometry, contrast sensitivity, and ocular higher-order aberrations [8,9]. However, only one previous report documented the posterior corneal impact after suspending RGP contact lens wear for 7 days [10], so the effects on the posterior cornea immediately after suspending RGP contact lens wear are not clear.

The development of corneal tomography techniques, such as anterior segment optical coherence tomography (AS-OCT), has allowed a more detailed and precise analysis of the anterior and posterior corneal configurations. The importance of posterior surface alterations in early keratoconus screening has been shown in several studies [11-13], but no consensus yet exists in the literature regarding the corneal stability in the posterior surface after removal of RGP contact lenses. Nevertheless, a better understanding of the changes in the shape of posterior cornea after RGP contact lens removal could be useful for practitioners who detect keratoconus and determine the indication of corneal cross-linking in patients wearing RGP contact lenses.

The aim of the present study was to evaluate the impact of RGP contact lens removal on the anterior and posterior corneas in eyes with keratoconus.

## Material and Methods

### Patient Inclusion

Eight patients with previously diagnosed keratoconus participated in the investigation. All subjects were recruited from the keratoconus clinic at the Kyoto Prefectural University of Medicine, Kyoto, Japan. All participants habitually wore RGP contact lenses. The patients were 34.3 ± 15.3 years old (mean ± standard deviation (SD); range 19–60 years) and included 5 men and 3 women. The research protocol followed the tenets of Declaration of Helsinki and was approved by the Kyoto Prefectural University of Medicine, Clinical Research Review Board, Kyoto, Japan (Approval # 1235), which is an independent organization for approval of ethical issues. Written informed consent was obtained from all subjects prior to their participation in the study. All patients had been diagnosed with keratoconus based on slit-lamp examinations that showed one or more of the following clinical findings: corneal Fleischer ring, Vogt’s striae, corneal thinning, and corneal protrusion at the apex and additionally if AS-OCT revealed a keratoconic appearance, such as focal or inferior steepening, a bow-tie pattern, or skewed axes [14]. Exclusion criteria were the following: previous ocular surgery, corneal scarring, trauma, acute corneal hydrops, or a history of other ocular disease beside refractive errors. Patients were instructed to wear their own RGP contact lenses more than 8 hours a day for up to 4 weeks before the assessment.

### Imaging Methods

Corneal topographic examinations using AS-OCT (SS-1000 CASIA; Tomey Corporation, Nagoya, Japan) were performed in all patients at 1, 5, 10, 20, and 60 minutes after RGP contact lens removal. The changes in the indices of the AS-OCT between 1 minute and 60 minutes after lens removal were then analyzed. The measurements obtained at 5, 10, and 20 minutes after RGP contact lens removal were used as reference data. All topographic examinations were performed by the same examiner (M.I.). We previously confirmed that data from AS-OCT have high reproducibility and repeatability [15]. The AS-OCT used in the present study was a swept-source OCT with a center wavelength of 1,310 nm and scan rate of 30,000 A-scans per second. A total of sixteen cross-sectional images, consisting of 512 A-scans, was obtained for 0.34 seconds during 1 measurement to assess the corneal topography. The AS-OCT measurements included anterior and posterior mean steep keratometry (Ks), mean flat keratometry (Kf), central corneal thickness (CCT) at the apex, corneal thickness at the thinnest point (CTmin), anterior or posterior best-fit sphere (BFS), elevation values, corneal surface areas, and the anterior-posterior ratio of the corneal surface area (As/Ps). The BFS was generated using built-in AS-OCT software with a float option within the central 5.0 mm of the cornea and used as a reference surface. Anterior or posterior elevations were measured as the maximum value above the BFS in the central 5.0 mm of the anterior or posterior cornea. The anterior or posterior corneal surfaces were calculated within the central 5.0 mm of the cornea based on the anterior or posterior elevation maps, as previously reported [16]. Briefly, we used the formula S = 2×PI×R (R-√R2-(D/2)2, in which S = surface area, PI = ratio of a circle’s circumference (3.14), R = curvature, and D = corneal diameter, at each measurement point based on the elevation map calculated with AS-OCT. The As/Ps was calculated as follows: As/Ps = anterior corneal surface area / posterior corneal surface area. Each examined eye was required to have a corneal map with at least 9.0 mm of corneal coverage and no extrapolated data by AS-OCT.

### Contact Lens Materials

A slit-lamp examination and sodium fluorescein were used to assess the fitting of the subject’s habitual RGP contact lens. All contact lenses used in this study were Sun Contact Lens Mild II (Sun-contact lens, Kyoto Japan), manufactured from dextran-methylmethacrylate copolymer material and prescribed by fitting with an apical-touch approach. A representative example of the RGP contact lens fluorescein fitting pattern is shown in Fig 1.

**Fig 1.**
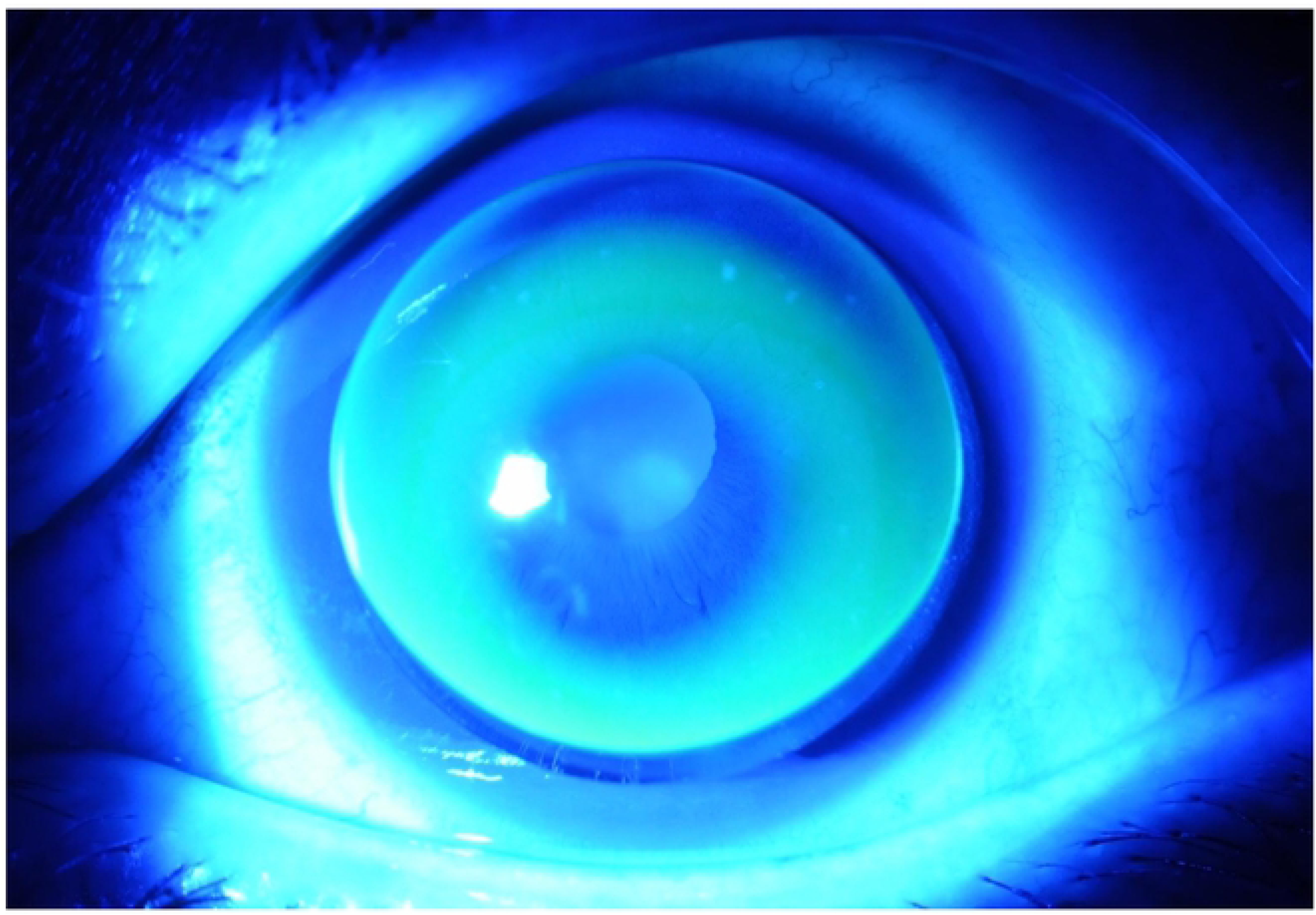
Fluorescein fitting patterns for rigid gas permeable contact lenses. (Case 3) Moderate keratoconus showed an apical touch lens fitting.

### Statistical Analyses

Statistical analyses were conducted using R version 3.1.0 (The R Foundation) statistical software, with the data presented as mean ± SD where applicable. The data were not normally distributed; therefore, the Wilcoxon signed-rank test was used to compare each parameter between the data at 1 minute and 60 minutes after RGP contact lens removal. Differences in the change rate of indices, the change between the data at 1 minute and 60 minutes after RGP contact lens removal, were compared using the Dunnett test. A *P*-value less than 0.05 was considered statistically significant.

## Results

Of the eight corneas analyzed, the mean Ks, Kf, and CCT at initial examination were 51.88 D (range 42.84–60.94 D), 48.85 D (range: 42.16–56.54 D) and 428.13 μm (range: 224–558 μm). The Amsler-Krumeich classification was used to classify the severity of keratoconic eyes into 4 groups: Grade 1 (2 eyes), Grade 2 (2 eyes), Grade 3 (2 eyes), and Grade 4 (1 eyes). A summary of the 8 patents with keratoconus at the initial examination is shown in Table 1.

**Table 1.** Demographic and topographic data and Amsler-Krumeich classification of the corneas in eight patients with keratoconus at initial examination.

| Case | Age<br>(years) | Sex | Ks<br>(D) | Kf<br>(D) | CCT<br>(μm) | A-K<br>grade |
| --- | --- | --- | --- | --- | --- | --- |
| 1 | 60 | F | 60.94 | 56.54 | 224 | IV |
| 2 | 54 | M | 53.06 | 50.98 | 432 | II |
| 3 | 21 | F | 55.44 | 54.28 | 412 | III |
| 4 | 38 | F | 44.96 | 42.72 | 461 | I |
| 5 | 19 | M | 52.27 | 47.13 | 465 | II |
| 6 | 27 | M | 43.57 | 42.65 | 488 | I |
| 7 | 25 | M | 42.84 | 42.16 | 580 | I |
| 8 | 30 | M | 61.96 | 54.3 | 363 | III |

Ks: Steep keratometry, Kf: Flat keratometry, CCTmin: Central corneal thickness, A-K: Amsler-Krumeich classification, F: Female, M: Male.

In all cases, both the anterior and posterior cornea showed steepening immediately after suspending the wearing of the RGP contact lenses, which were fitted using an apical touch fitting approach, as shown in Fig 1. Statistically significant steepening was noted for topographic parameters (BFS and elevation values) of both the posterior and anterior corneal surfaces in the first 60 minutes after removal of the RGP contact lens. The anterior and posterior corneal surface areas significantly increased, but the As/Ps ratio showed no significant difference in the first hour after RGP contact lens removal. (Table 2) Representative examples of AS-OCT images performed at 1 minute and 60 minutes after RGP lens removal are shown in Fig 2.

**Table 2.** Summary of topographic changes in the anterior and posterior corneal surfaces after RGP contact lens removal.

| Index | Time after lens removal |  |  |  |  | 1min vs. 60 min |
| --- | --- | --- | --- | --- | --- | --- |
|  | 1 min | 5 min | 10 min | 20 min | 60 min | (P-value) |
| aBFS | 6.6238 | 6.5688 | 6.5363 | 6.5413 | 6.4488 | < 0.01 |
| pBFS | 4.9363 | 4.8913 | 4.8800 | 4.8475 | 4.8463 | < 0.01 |
| AE | 47.5250 | 48.8125 | 49.3125 | 51.1625 | 52.4250 | < 0.01 |
| PE | 115.7125 | 115.4875 | 115.9875 | 115.8125 | 116.7375 | < 0.01 |
| CTmin | 380.0000 | 373.0000 | 368.1250 | 376.6250 | 375.8750 | < 0.01 |
| As | 20.3635 | 20.4129 | 20.4241 | 20.4290 | 20.4421 | < 0.01 |
| Ps | 21.0035 | 21.0565 | 21.0749 | 21.0791 | 21.0901 | < 0.01 |
| As/Ps | 0.9698 | 0.9698 | 0.9695 | 0.9696 | 0.9696 | 0.105 |

aBFS: anterior best-fit sphere, pBFS: posterior best-fit sphere, AE: anterior elevation values, PE: posterior elevation values, CTmin: corneal thickness at the thinnest point, As: Anterior corneal surface area, Ps: posterior corneal surface area, As/Ps: the anterior-posterior ratio of the corneal surface area, min: minute, vs: versus

**Fig 2.**
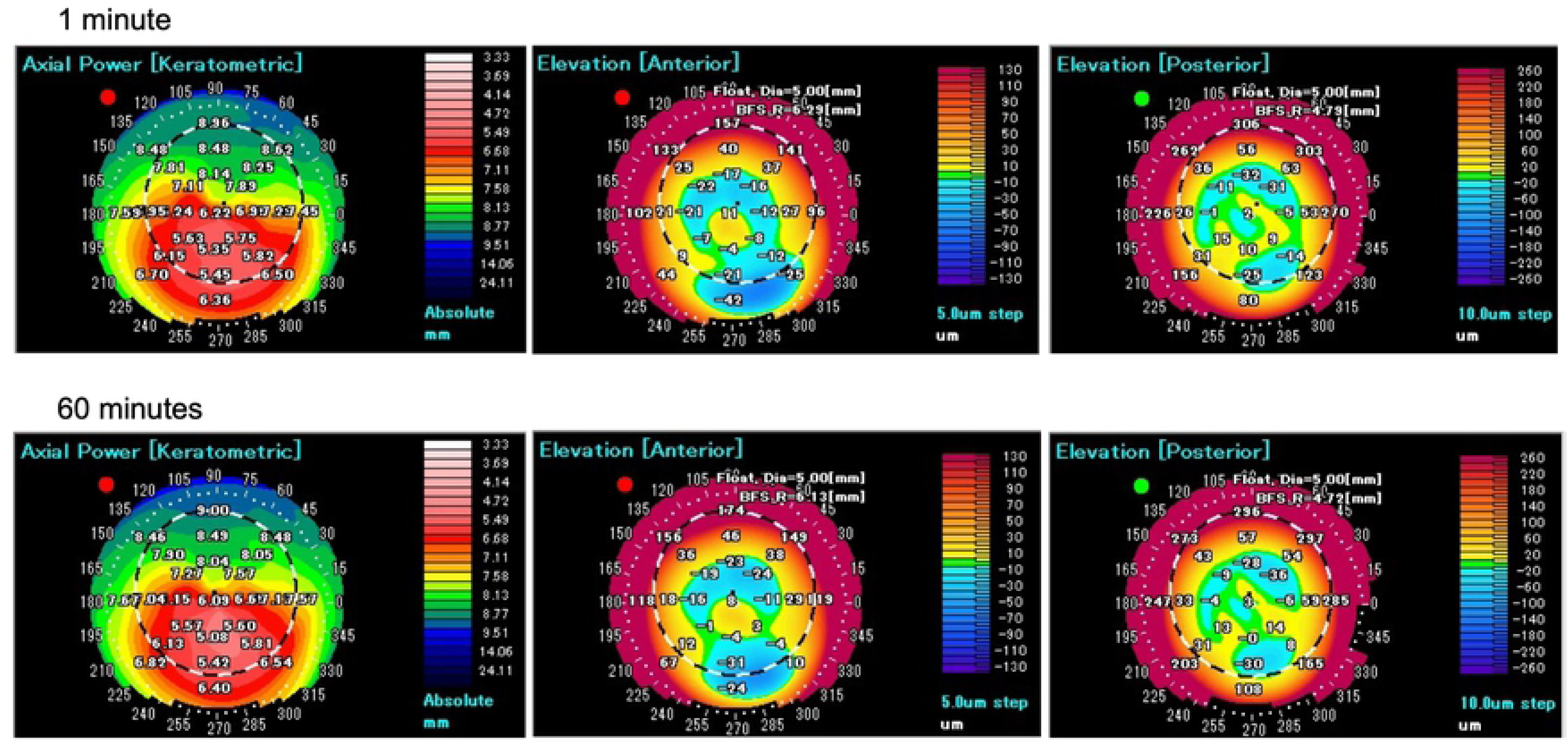
Anterior segment optical coherence tomography images performed at 1 minute and 60 minutes after removal of the rigid gas permeable contact lens. (Case 2) Anterior Axial map (left-side columns), and anterior and posterior elevation map (central and right columns) showed steepening between 1 minute (upper) and 60 minutes (lower) after removal of the rigid gas permeable contact lens.

## Discussion

The results from this study confirmed significant effects on both the anterior and posterior cornea in patients with keratoconus in the first 60 minutes after removal of RGP contact lenses. Previous studies reported no significant changes in the posterior cornea after suspending RGP contact lens wear for 7 days; [10] however, to the best of our knowledge, the present study is the first to report an initial impact on the posterior corneal surface after discontinuing the wearing of RGP contact lenses.

The findings of the present study have established that both the anterior and posterior surfaces of the cornea showed steepening at 60 minutes after removal of the RGP contact lenses. This steepening is likely to represent a recovery from the “molding” [17] effect of RGP contact lens wear. However, Jinabhai et al. reported no significant differences in the posterior cornea after suspending RGP contact lens wear, although a change was noted in the anterior cornea [10]. This discrepancy could be explained by the difference in the fitting approach used for the RGP contact lens. In addition to a longer duration of lens wear [18, 19], the fitting approach for the RGP contact lens has been reported to influence the topographic changes induced by RGP contact lens wear. For example, Romero et al. showed that the apical touch fitting approach was associated with a greater corneal flattening when compared to three-point-touch lens wear [7]. In the present study, all lenses were prescribed with apical touch, and this fitting approach may induce significant flattening that results in subsequent recovery of the posterior corneal shape.

The results from the present study indicated significant changes in the anterior and posterior BFS, elevation values, and surface areas in the first 60 minutes after removal of the RGP contact lens. These results suggest that, in the first hour after suspending the wearing of RGP lenses fitted with apical touch, the keratoconus screening index derived from the posterior corneal surface, as well as from the anterior corneal surface, may be affected by removal of the RGP contact lens. We should take this steepening in the cornea into consideration to prevent underestimation when evaluating the corneal topography after removal of RGP contact lenses.

We previously reported that AS-OCT could provide a measurement of the corneal surface area, obtained by multiplying the corneal curvature [16], and that the As/Ps ratio may reflect the precise corneal shape as a 3D structure [15]. The results from the present study indicated a significant increase in the anterior and posterior surface areas in the first 60 minutes after RGP contact lens removal. These findings suggested the occurrence of a steepening of the cornea in both the anterior and posterior surfaces after suspending RGP contact lens wear, which was consistent with the effects on the BFS and elevation values. Conversely, the As/Ps ratio did not show any statistically significant change in the first hour after RGP contact lens removal, thereby suggesting, theoretically, that the As/Ps index minimizes the impact due to certain changes in the anterior and posterior surface area when RGP contact lens wear is discontinued. However, the complete picture of the overall corneal changes after RGP contact lens removal still remains unclear. Our findings indicated that the As/Ps ratio in the keratoconic cornea is unchanged, suggesting that the anterior and posterior corneal surfaces undergo a similar shift after RGP lens removal.

A possible limitation is that the study focused on the effects occurring in the first 1 hour after removal of the RGP contact lenses. Previous reports have indicated that recovery of normal stability of the corneal topography takes 1 to 21 weeks in normal subjects after suspending RGP contact lens wear [20-23]. A longer study might be necessary to determine how long these changes persist in the keratoconic cornea.

In conclusion, our findings show that patients with keratoconus experienced significant changes in both the anterior and posterior cornea areas in the first 60 minutes after removal of RGP contact lenses. Further investigations into the development of new screening indexes for keratoconus that consider the impact of RGP contact lens wear on corneal shape would be clinically useful for clinicians who diagnose keratoconus and for determining whether corneal cross-linking is indicated.

## ACKNOWLEDGMENTS

The authors wish to thank Dr. Yoko Hyakutake for insightful suggestions.

## Author Contributions

**Conceptualization:** Motohiro Itoi, Koji Kitazawa.

**Data curation:** Motohiro Itoi

**Formal analysis:** Motohiro Itoi, Koji Kitazawa

**Investigation:** Motohiro Itoi, Koji Kitazawa

**Methodology:** Motohiro Itoi, Koji Kitazawa

**Project administration:** Motohiro Itoi, Koji Kitazawa

**Resources:** Motohiro Itoi, Koji Kitazawa, Hiosayo Higashihara, Chie Sotozono

**Software:** Morohiro Itoi

**Supervision:** Koji Kitazawa, Hisayo Higashihara, Chie Sotozono

**Validation:** Motohiro Itoi, Koji Kitazawa

**Visualization:** Motohiro Itoi, Koji Kitazawa

**Writing – original draft:** Motohiro Itoi, Koji Kitazawa

**Writing – review & editing:** Koji Kitazawa, Hisayo Higashihara, Chie Sotozono

